# Broad intraspecies killing activity in *Pseudomonas syringae* due to the combinatorial action of LPS-interacting bacteriocins

**DOI:** 10.1101/2023.09.27.559845

**Authors:** Savannah L. Weaver, Emma Casamassima, Anh Evy Nguyen, David A. Baltrus

**Affiliations:** School of Molecular and Cellular Biology, University of Arizona, Tucson AZ, USA; School of Plant Sciences, University of Arizona, Tucson AZ, USA; School of Animal and Comparative Biomedical Sciences, University of Arizona, Tucson AZ, USA

## Abstract

Bacteriocins are a diverse group of highly specific antimicrobials produced by bacteria, thought to mainly target and kill strains that are closely related to and which therefore potentially compete in the same niche space as producer cells. Single strains can produce more than one type of bacteriocin, with each usually having differing modes of action and receptors for binding, and with strain specificity for each independent bacteriocin due to the requirement for these molecules to bind to receptors in target cells prior to carrying out antibacterial functions. Here we show that *Pseudomonas syringae* pv. aptata DSM50252 (*Ptt*) displays broad intraspecific killing activity due to combinatorial and non-overlapping activities of phage derived bacteriocins (referred to as tailocins) as well as a prophage encoded lectin-like bacteriocin (aptatacin L1). These results highlight how single strains can maintain broad killing activity against a variety of potential competitors by targeting multiple conformations of a shared receptor, and provide additional evidence that tailocins and aptatacin L1 both utilize rhamnose moieties in the LPS as potential receptors for binding.

## INTRODUCTION

Microbial community structure can be driven by antagonistic interactions within and between strains and species, leading to increased interest in mining the molecules behind such interactions as next generation antimicrobials (1, 2). *Pseudomonas syringae* pv. aptata DSM50252 (*Ptt*) was demonstrated to possess the broadest intraspecies killing spectrum compared to other *P. syringae* strains in a relatively large screen for activity of phage derived bacteriocins (referred to hereafter as tailocins, (3)). A better understanding of the genetic factors enabling *Ptt* to maintain extensive intraspecies antibacterial activity could provide insights into how to develop optimized antimicrobials and better deploy antimicrobials in effective combinations (4–7), and we therefore sought to identify the underlying basis for this broad-spectrum activity.

Bacteriocins are proteinaceous antibacterial compounds produced by bacteria that typically have activity against closely related strains, either by interrupting processes within the cell or disrupting membrane integrity (8–11). Production of these molecules is typically thought to be coregulated with the SOS response and upregulated in response to DNA damage, as they are often repressed by the action of LexA-like proteins or LexA itself (12–15). Bacteriocins have already shown promise as effective and highly specific treatment options to prevent or treat bacterial infections through application and when engineered into host genomes (6–8, 16). A deeper understanding of the modes of actions of multiple bacteriocins produced by single strains could therefore aid the development and optimization of future bacteriocin treatments across both clinical and agricultural settings.

Specificity of bacteriocin targeting is imparted by a requirement of these proteins to bind to receptors on target cells prior to carrying out antimicrobial functions, and bacteriocin classes can therefore be separated by their molecular composition, size, and modes of function (8). Diffusible bacteriocins, such as colicins and microcins, are relatively small molecules that diffuse or are translocated across the cell membrane of target bacteria to enter the cytoplasm, where antimicrobial activity is often due to DNAse or RNAse activity (17–19). Tailocins are relatively large phage-derived particles that are thought to bind to the receptors (like the lipopolysaccharide, LPS) that decorate the outside of target bacteria, and which are thought to puncture the cell wall with their spike protein causing disruption of the membrane potential and therefore cell death (13, 20–22). Lectin-like bacteriocins produced by *Pseudomonas spp.* were previously thought to bind targeted D-mannose oligosaccharides, but have now been shown to utilize D-rhamnose of the LPS in a genus-specific manner (23, 24). The exact mechanism of action by lectin-like bacteriocins remains unknown, but studies have demonstrated that interactions with both the LPS and the outer membrane protein BamA are required for lethal activity (24).

We previously used polyethylene glycol (PEG) precipitation to screen for tailocin-like killing activity across diverse strains of the phytopathogen *P. syringae*, and found *Ptt* maintained the broadest target range according to these assays (3). By using PEG to precipitate high-molecular weight compounds, apatacin L1 theoretically should not be left in the precipitated supernatant since lectin-like bacteriocins consist of a single protein versus tailocins that form large protein complexes (25, 26). We follow up on these initial results by demonstrating that only a subset of witnessed killing activity post PEG precipitation for this strain can be attributed to tailocins. We further show that the remaining activity is due to the presence the lectin-like bacteriocin, aptatacin L1, which shares similarity with the putidacin L1 described by McCaughey et al. Together, these results characterize and clarify the underlying molecular and genetic basis of broad-spectrum killing activity by *Ptt* against other *P. syringae* strains.

## RESULTS

### An Improved, Nearly Complete, Genome Sequence for *P. syringae* pv. *aptata* DSM50252

As a first step towards fully characterizing bacteriocin production by *Ptt*, we generated a new, nearly complete, genome assembly for this strain. This whole genome shotgun project has been deposited at DDBJ/ENA/Genbank under accession AEAN00000000 and was sequenced independently from the original assembly (27). The version described in this paper is version AEAN02000000. This genome assembly is composed of one circular chromosome and one circular plasmid (pPttDSM50252-1; 53,430bp). Curiously, despite the chromosome being circularized by the unicycler pipeline, one additional 695,685bp linear contig was produced during assembly. Given the content of this contig (which includes a variety of well known “housekeeping” genes for *Pseudomonas* like *gacA* which should be found within the *Ptt* chromosome, we assume that circularization was a misassembly error and therefore include this additional contig as part of the genome assembly. The total size of all three contigs and therefore the genome is 5,918,862bp.

### *Ptt* shows a variable and differential killing phenotype in overlay assays

Our original results demonstrated that *Ptt* possesses the broadest spectrum of killing activity compared to other *P. syringae* strains that were screened (3). As part of the screen, we were able to classify *P. syringae* strains into two broad sensitivity classes (A and B) according to their interactions with PEG precipitated supernatants produced by diverse *P. syringae* strains. Unlike many other strains, PEG precipitated supernatants from *Ptt* maintained activity against many strains from both tailocin sensitivity classes. Reinspection of overlays behind these results highlighted an additional phenotype of interest relevant to the broad killing spectrum of *Ptt* in that the zone of killing for *Ptt* preparations was visibly larger against sensitivity class A strains than against class B strains (Figs. 1A and 1B). Killing zones were also visibly larger for preparations from *Ptt* against class A strains when compared to other tailocins known to target class A strains (Figs. 1A and 1B). As a follow up experiment, we retested the sizes of killing zones when class A and B strains were treated with unselected supernatants from tailocin induced cultures of *Ptt* and USA011. Using *Psy*B728a and *P. syringae* UB303 as representative target strains for tailocin sensitivity class A and B strains, respectively, as shown in Fig. 1 our new data clearly demonstrates that the killing zone from *Ptt* supernatants is much larger against tailocin sensitivity class A strains than tailocins from strain USA011 against these same targets. The zone of inhibition is also larger than killing zones generated by *Ptt* supernatants against tailocin sensitivity class B strains. We further find that the area of the PEG prepped supernatant killing zone is not significantly smaller compared to raw supernatant (Fig. 1B), although variability in the assay may be obscuring an underlying trend. These results indicate that PEG precipitation does not cleanly select out a subset of the killing activity from these supernatants, unlike other strains where diffusible bacteriocins are eliminated by this treatment (26).

**Figure 1.**
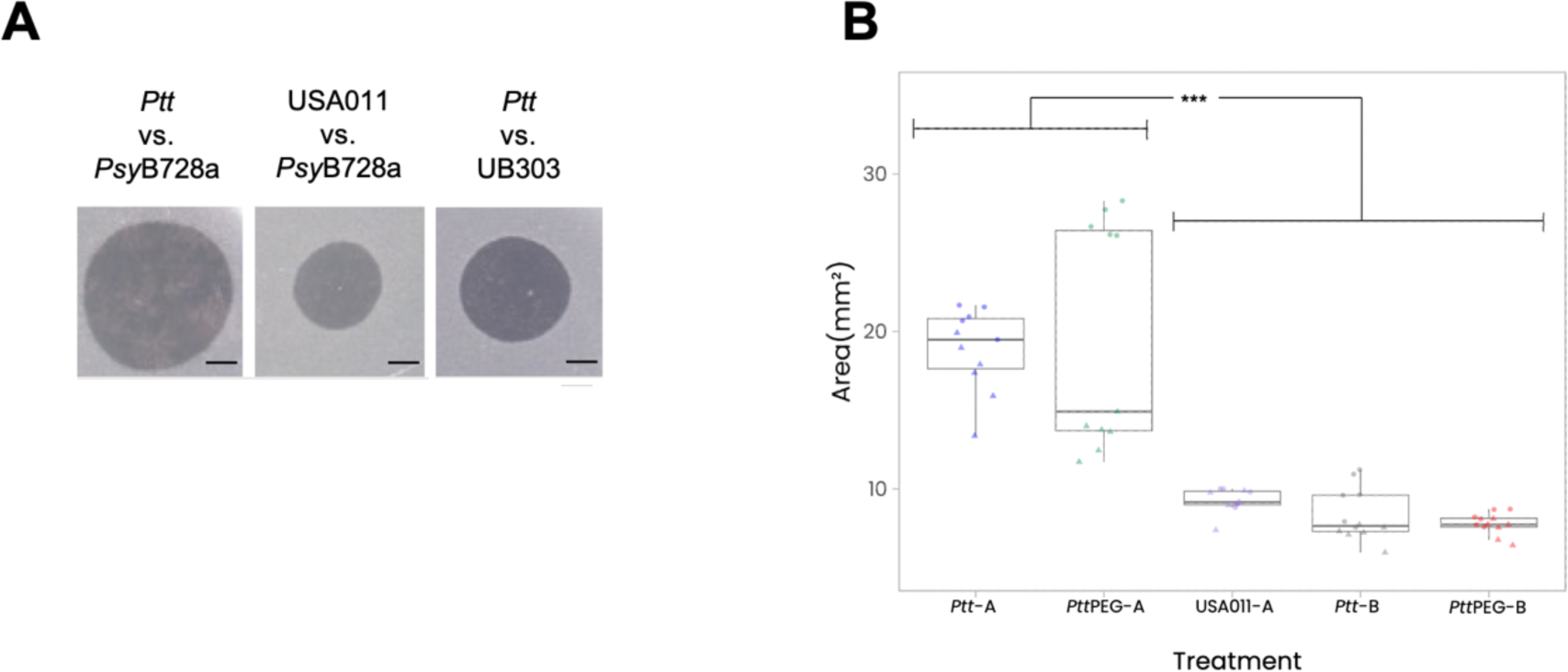
*Ptt* supernatants produce a relatively large killing zone against sensitivity class A strains. A) Overlays comparing killing zone size differences with a 1mm scale bar for comparison. From left to right, *Ptt* treatment shows a larger than normal killing zone against sensitivity class A strain B728a as the overlay strain, USA011 treatment shows a normal size killing zone against sensitivity class A strain B728a as the overlay strain, and *Ptt* treatment shows a normal size killing sone against sensitivity class B strain UB303 as the overlay strain. B) Data from overlay assays comparing killing zone sizes: *Ptt* and *Ptt*PEG on sensitivity class A strain B728a overlays, USA011 on sensitivity class A strain B728a overlays, and *Ptt* and *Ptt*PEG on sensitivity class B strain UB303 overlays. *Ptt* and *Ptt*PEG against sensitivity class A strain B728a showed significantly larger killing zones compared to USA011 against B728a and *Ptt* and *Ptt*PEG against sensitivity class B strain UB303 (Tukey test ***p<0.0001). Assays were performed two independent times, with three replicates each. Scans underlying images can be found at raw data and code to reconstruct Figure 1B can be found at https://doi.org/10.5281/zenodo.8384752

### Killing activity against tailocin sensitivity class A strains is eliminated in a tailocin knockout strain

In previous studies, creation of a double knockout mutant through the deletion of the receptor binding protein (Rbp) and adjacent chaperone (Ch) from the tailocin locus was sufficient to abolish killing activity in a particular strain (3, 7, 28). To investigate tailocin killing activity in *Ptt*, we created a similar a Rbp/Ch double knockout mutant and tested for killing activity against strains from tailocin sensitivity classes A and B. Instead of eliminating all killing activity from supernatants after PEG precipitation, as one would predict if the activity was solely due to tailocins, we observed that the Rbp/Ch knockout (*Ptt*Δ*rbp/ch*) abolished killing against all tailocin sensitivity class B strains but maintained activity against all sensitivity class A strains (Fig. 2).

**Figure 2.**
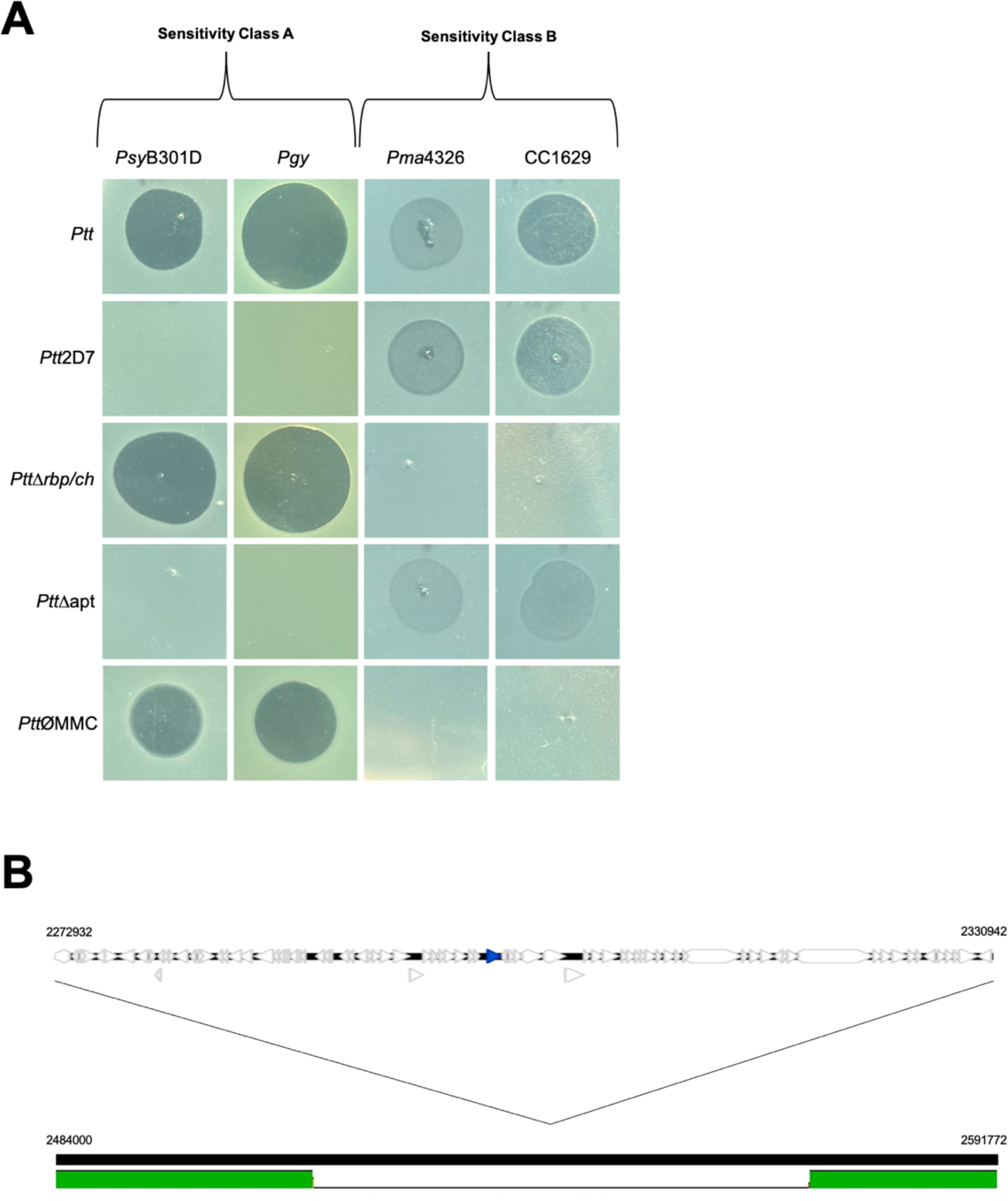
*Ptt* killing activity against different sensitivity classes due to combinatorial activity of two different bacteriocins. A) Rows are bacteriocin treatment conditions and columns are target strains sectioned out into sensitivity classes A and B. *Ptt* shows killing zones across both sensitivity classes, with killing zone sizes being visibly smaller against class B. A transposon mutant, *Ptt*2D7, is missing a prophage region, and subsequently aptatacin L1, in the genome and shows abolished activity against class A strains. The tailocin knockout mutant, *PttΔrbp/ch*, shows abolished activity against class B strains. The aptatacin L1 knockout mutant, *PttΔapt,* shows abolished activity against sensitivity class A. Lastly, *Ptt* wildtype cultures not induced by MMC only show activity against class A strains. These results, together, indicate that *Ptt* tailocins target and kill sensitivity class A strains and aptatacin L1 targets and kills sensitivity class B strains. B) Genome alignment (bottom) of *Ptt* wildtype vs. transposon mutant *Ptt*2D7 showing missing prophage region indicated by the white region in between the two outside green sections. A visualization of the prophage genes is included above with the *aptatacin L1* gene colored blue. Scans underlying cropped images can be found at https://doi.org/10.6084/m9.figshare.24208449

### Non-tailocin antimicrobial activity is due to a phage derived lectin-like bacteriocin

Given that killing activity was maintained against tailocin sensitivity class A *P. syringae* strains in the Rbp/Ch deletion mutant, we carried out transposon mutagenesis to identify genes responsible for remaining antimicrobial activity. We were successfully able to isolate multiple mutants with independent transposon insertions each of which had lost the antimicrobial activity against tailocin class A strain *Psy*B728a. Focusing on transposon mutant *Ptt*2D7, we further and found that killing activity in this mutant was abolished against other strains designated as tailocin sensitivity class A but antagonistic activity was maintained against tailocin sensitivity class B strains (Fig. 2).

However, investigation of transposon insertion positions in *Ptt*2D7 as well as two additional and independent transposon strains where activity against class A strains was abolished strongly suggested that transposon insertions were not causative of loss of antimicrobial activity as each of the insertions was in unrelated genes predicted disrupt pathways involved in metabolism and growth. Specifically, the transposons were found to have disrupted *hopAC* (PSYAP_002225), an ABC transporter (PSYAP_017405), and *mksF* (PSYAP_003975), in transposon strains 2D7, 4E3, and 5D10 respectively. Further investigation of genome assemblies for these three strains showed that, in addition to possessing independent transposon insertions, each had also lost a prophage compared to the original *Ptt* genome (Fig. 2B). Evaluation of genes encoded by this prophage using BagelIV highlighted the presence within the phage genome of an ORF encoding a potential lectin like bacteriocin previously identified in strain *Ptt* by McCaughy et al. and which we will refer to as aptatacin L1. (23, 29, 30).

### *Ptt* tailocin targets sensitivity class B strains, while aptatacin L1 targets sensitivity class A

To demonstrate that aptatacin L1 was responsible for killing activity maintained in the *Ptt* Rbp/chaperone mutant against tailocin sensitivity class A strains, we deleted the aptatacin L1 gene (*apt)* from the *Ptt* genome and evaluated killing activity of this mutant. We find that killing activity is abolished when supernatants from *Ptt*Δ*apt* are tested against multiple strains classified as sensitivity class A, but is maintained against all strains classified as sensitivity class B (Fig. 2). In parallel, we cloned the *apt* gene under control of a *lac* promoter into pET-28a(+) and measured the killing activity arising from supernatant from *E. coli* cultures induced by IPTG. As one can see in Figure 3, production of aptatacin L1 by *E. coli* is sufficient to enable killing against *P. syringae* class A strains (Fig. 3B). Therefore, it is clear that killing activity of *Ptt* against sensitivity class A strains is due to the production of aptatacin L1.

**Figure 3.**
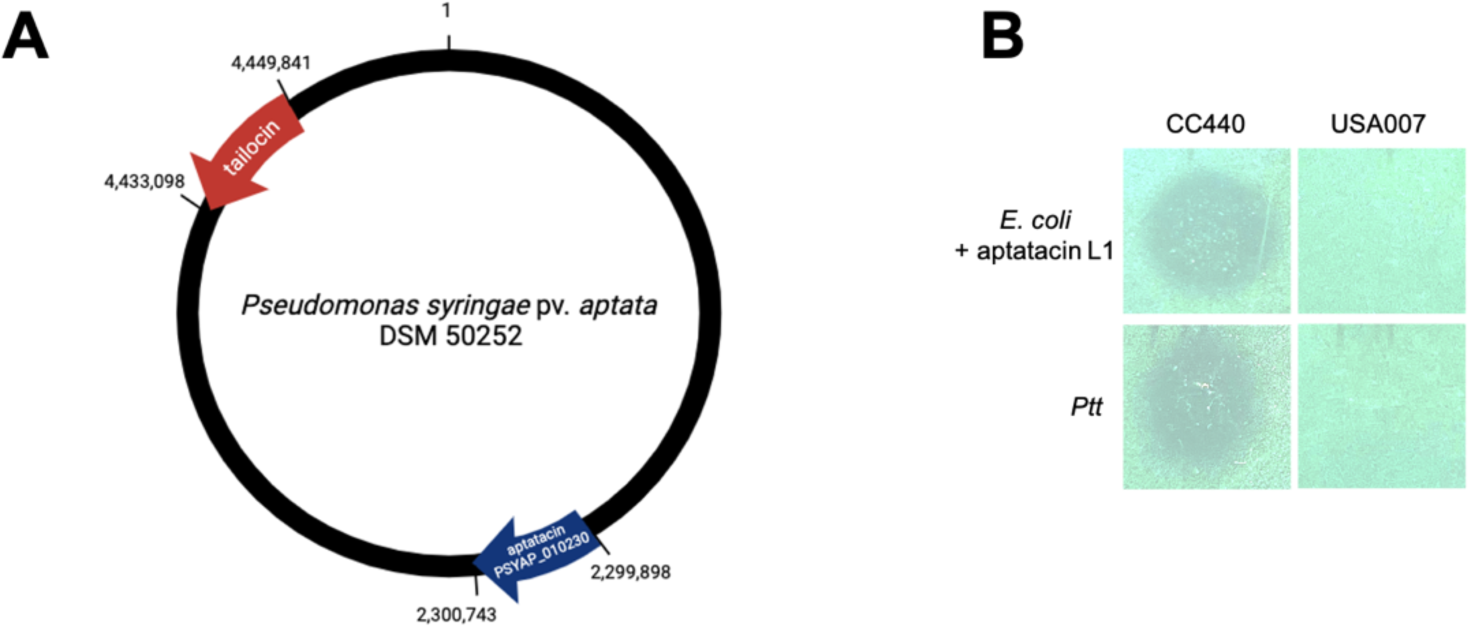
Production of aptatacin L1 is differentially regulated from tailocins. A) Genome visualization of *Ptt* strain DSM50252 showing the distinct locations of the tailocin locus and *aptatacin L1* gene. The argument can be made that these two bacteriocins are under two different regulatory pathways given the distance between the loci. B) Induction of aptatacin L1 in *E. coli* strain BL21 shows activity against sensitivity class A strain CC440 that *Ptt* also shows activity against. There is no activity shown against sensitivity class A strain USA007, which is not a target of *Ptt*.

### Low level of induction of aptatacin L1 under normal laboratory conditions, with further induction by mitomycin C

*P. syringae* tailocin production is thought to be induced by DNA damage, as tailocin biosynthesis pathways are repressed by a *lexA*-like regulator (referred to as *ptrR* in *P. aeruginosa* but with a similar protein present in *P. syringae* (12, 14, 31)). Since the *apt* gene is present in a prophage in a different part of the chromosome than the tailocin locus (Fig. 3A), we tested for induction of this lectin-like bacteriocin by DNA damage in response to mitomycin C (MMC). We find that *Ptt* shows killing activity against sensitivity class A strains with and without MMC induction, although the activity is lowered in the absence of MMC. In contrast, tailocin-dependent killing activity by *Ptt* against sensitivity class B strains is seen only after induction by MMC (Fig. 2). These results therefore indicate that *apt* is expressed and aptatacin L1 appears to be produced at a low level regardless of DNA damage under normal laboratory conditions, but also suggests that the locus encoding aptatacin L1 can be additionally upregulated by the presence of DNA damage.

## DISCUSSION

Bacteriocins possess great potential for use as highly efficient strain-specific antibiotics to replace, or work alongside, broad-spectrum antibiotics for the prevention and treatment of bacterial infections across clinical and agricultural settings (8). Before deployment in such cases, we must first gain a deep understanding of the underlying regulatory mechanisms by which they are produced and released, how target host strains are sensed by the bacteriocins, the genetic basis for target strain specificity, as well as the mechanism of action of different subsets of bacteriocins. Towards these goals, we sought to investigate the genetic basis underlying exceptionally broad intra-species killing by bacteriocins produced by *Pseudomonas syringae* pv. aptata DSM50252 (*Ptt)*.

Despite an extensive amount of research identifying and characterizing bacteriocins across bacteria, and recent insights into evolutionary costs and benefits of employing different types of bacteriocins, much less is known about how evolutionary pressures structure bacteriocin composition within single strains (32–34). Given that our previous publication (3) screened for bacteriocin activity from supernatants that had undergone PEG precipitation (which we assumed cleanly selects tailocins away from other diffusible bacteriocins), our initial hypothesis was that the ability of *Ptt* to target strains from multiple tailocin sensitivity classes meant that tailocins from *Ptt* possessed unique capabilities compared to those from other strains. However, further inspection of overlay data showed that killing zones from *Ptt* PEG-precipitated supernatants were larger against tailocin sensitivity class A strains than against class B strains even when the same volume of supernatant was applied, suggesting that killing activity was not phenotypically homogenous against all other *P. syringae* strains. As relayed above, our follow up results demonstrate both that non-tailocin bacteriocins can be pulled down during PEG precipitation despite their smaller size but also highlight that different types of bacteriocins can have similar or analogous receptors in target strains.

There is strong genetic evidence that tailocins produced by various Pseudomonads interact with sugar chains that compose the O-antigen of the LPS to mediate antimicrobial activity (35–38). Moreover, multiple studies strongly suggests that *Ptt* and other strains classified as tailocin sensitivity class A appear to maintain an LPS O-antigen region consisting of D-rhamnose while sensitivity class B strains maintain an O-antigen that consists of L-rhamnose (37, 39, 40). Data reported here further supports this hypothesis, as an Rbp/chap tailocin deletion strain fails to kill sensitivity class B *P. syringae* strains and thus it is likely that the *Ptt* tailocin targets strains with L-rhamnose moieties in their LPS to complement the activity of aptatacin L1 against D-rhamnose moieties as described below.

Lectin-like bacteriocins, such as aptatacin L1, have been shown to interact with D-rhamnose of the LPS and likely bind it (23), and thus potentially are able to target tailocin sensitivity class A strains, but also ultimately mediate killing activity through further interactions with the BamA protein (24). Since *Ptt* is classified as a tailocin sensitivity class A strain, and therefore likely possesses a D-rhamnose based O-antigen, we predict that aptatacin L1 should be able to bind to the LPS of *Ptt.* However, inspection of sequence variation in *bamA* from the *Ptt* genome highlights that this strain likely avoids killing by aptatacin L1 despite possessing D-rhamnose because its version of BamA possesses a motif that is immune to lectin-like bacteriocin killing (24). Regardless of the status of BamA across other genomes of *P. syringae*, sensitivity class B strains fail to be killed by aptatacin L1 because this bacteriocin cannot bind to the L-rhamnose based LPS of these strains and indeed we witnessed no killing activity against class B strains from our tailocin Rbp/chap deletion strain. Therefore, our results demonstrate that the previously reported broad killing spectrum of *Ptt* PEG precipitated supernatants is due to the combination of activities of two different types of bacteriocins: a tailocin which kills strains with L-rhamnose based O-antigen (sensitivity class B) and aptatacin L1 which kills strains with D-rhamnose based O-antigen and which also possess *bamA* alleles that are sensitive to lectin-like bacteriocins. Furthermore, since other tailocins produced by *P. syringae* are predicted to interact with D-rhamnose, it remains an open question whether aptatacin L1 and D-rhamnose targeting tailocins interact with the O-antigen of the LPS in similar ways.

Phages are known to encode pathways to produce antimicrobial compounds which likely benefit both the host cell and the phage by allowing bacterial hosts to outcompete other cells (41) Resistance to many bacteriocins is usually provided by resistance genes that are coproduced together with the bacteriocin, rendering the producer cells inherently immune to these toxins. It is therefore straightforward to imagine how such bacteriocin-resistance gene modules can be acquired by phage and plasmids, as their modular setup renders activity of the bacteriocin largely independent of plasmid and phage functions. The presence of a gene encoding aptatacin L1 within a prophage, as is found in *Ptt* as well as for other lectin-like bacteriocins, presents a slightly more confounding situation because there doesn’t appear to be other LPS-modification genes present in this phage locus in *Ptt.* Therefore, it is possible that the presence of lectin-like bacteriocins potentially limits host range of the phage and ultimately affects phage evolutionary dynamics because new potential hosts for the phage must not only have a receptor for the phage but must also possess an appropriate genetic background to be resistant to aptatacin L1.

Bacteriocins are typically produced by bacterial species as a competitive advantage against other bacterial species in a particular microbiome when there are stressors, such as limited resources (1, 2). Tailocin encoding operons are induced by DNA damage and tightly controlled at the transcriptional level, likely because all known tailocin loci also co-express lysozyme to enable bacteriocin release to the environment and thus expression of tailocin-coregulated operons is ultimately lethal. Given the phage-dependent location of aptatacin L1, we investigated the regulation of the *apt* gene. Although we saw that this gene was induced under DNA damage by MMC like tailocins, we witnessed aptatacin L1 production at lower and levels “normal” conditions of laboratory growth. Therefore, *Ptt* likely maintains the ability to kill strains of tailocin sensitivity class A in the absence of DNA damage, and this potentially allows for. This subtle change in regulation could also indicate that aptatacin L1 is effective in targeting other strains under slightly different conditions than tailocins.

As we observed previously, strain *Ptt* possessed the broadest target range when evaluated in the context of a relatively large and diverse screen for tailocin activity across *P. syringae* strains (3). Here, we demonstrate that this abnormally broad killing activity by *Ptt* can be attributed to the deployment of two classes of bacteriocins that act in a combinatorial way to target strains with multiple types of O-antigens in their LPS. Further investigation into the specific binding targets and regulatory pathways of these two types of bacteriocins will be useful to understand environmental conditions with which they are beneficial in nature but may also help to guide development of next generation strain-specific antimicrobials for use in agricultural conditions.

## METHODS

### Bacterial strains and growth conditions

*P. syringae* pv. *aptata* DSM50252 was originally isolated from sugar beet, and the isolate used for this manuscript displays resistance to rifampicin and was originally acquired from the Dangl lab collection (27). All strains and plasmids are listed in Table 1 and with descriptions of original isolation and modifications found in (42–44). Unless otherwise specified, all strains were grown in King’s B Media (KB) or on KB agar plates and antibiotics and other supplements were used in the following concentrations: kanamycin 25ng/uL, rifampicin 50ng/uL, nitrofurantoin 80ng/uL, tetracycline 10ng/uL, streptomycin 200ng/uL, diaminopimelic acid (DAP) 200ng/uL.

**Table 1.** Bacteria and Plasmids Used in This Study.

| Table 1. Bacteria and Plasmids Used in This Study |  |  |  |
| --- | --- | --- | --- |
| **Bacteria** |  |  |  |
| <u>Species and strain</u> | <u>Baltrus Lab</u> | <u>Origin and/or features</u> | <u>Source/citat</u> |
| *P. syringae* pv. *aptata* DSM50252 (*Ptt*) | DBL1559 | Dangl laboratory isolate | 27 |
| *P. syringae* USA011R | DBL1985 | Rifampicin resistant isolate of USA011 | 28, 43 |
| *P. syringae* UB303R | DBL1887 | Rifampicin resistant isolate of UB303 | This study |
| *P. syringae* pv. *syringae* B728a | DAB2 | Dangl laboratory isolate | 42 |
| *P. syringae* pv. *aptata* DSM50252 (*Ptt*) | DBL1944 | Tailocin receptor binding protein and chaperone deletion | This study |
| *P. syringae* pv. *aptata* DSM50252 (*Ptt*) | DBL1945 | *apt* deletion | This study |
| **Plasmids** |  |  |  |
| <u>Plasmid Name</u> |  | <u>Plasmid Features</u> | <u>Source/citat</u> |
| pDBL113 |  | pMTN1907 backbone for deletion of tailocin receptor binding protein and chaperone | This study |
| pDBL114 |  | pKM-18s backbone for deletion of *apt* | This study |
| pDBL115 |  | pET-28a(+) backbone for expression of *apt* | This study |

### Genome Sequencing and Assembly

The Baltrus Lab stock of *P. syringae* pv. *aptata* DSM50252 was originally acquired from the Dangl Lab stock in 2006 and frozen in 40% glycerol after two additional passages on KB agar plates. For each genomic DNA extraction used in the assembly reported here, a sample of this frozen stock was streaked onto KB agar plates, and a single colony was transferred to 2 ml of KB broth and grown overnight at 27°C in a shaking incubator at 220 rpm. Genomic DNA used for Illumina sequencing was isolated from a 2-ml overnight culture via the Promega (Madison, WI) Wizard kit following the manufacturer’s protocols and with the addition of RNAse. A separate genomic DNA extraction was performed, as above, for Illumina and Nanopore libraries. Sequencing of mutants followed the same protocols for DNA extraction.

DNA for the *Ptt* genome was sent to MiGS (Pittsburgh, PA) for Illumina sequencing following the standard workflow for library preparation and read trimming. As described in (45), this workflow uses an Illumina tagmentation kit for library generation, followed by sequencing on a NextSeq 550 instrument with 150-bp paired-end reads. Trimmomatic (46) was used for adaptor trimming using the default settings. This workflow generated a total of 1,331,568 paired reads and 358,000,888bp (∼60× coverage) of sequence. Raw Illumina reads have been uploaded to the SRA at accession SRR25551084.

DNA for the *Ptt* genome was sent to Plasmidsaurus (Eugene, OR) for Nanopore sequencing following the standard workflow for library preparation and read trimming. Samples were prepared using the Oxford Nanopore Technologies Ligation Sequencing Kit version SQK-LSK114, then sequenced on GridION 10.4.1 flowcells (FLO-MIN114) with the “Super accuracy” basecaller in MinKNOW using ont-guppy-for-promethion version 6.5.7.This workflow generated a total of 95,408 reads and 477,229,324bp (∼80× coverage) of sequence with a read N50 of 8750bp. Raw Nanopore reads have been uploaded to the SRA at accession SRR25551084.

For confirmation of the Rbp/ch deletion mutant and for identification of transposon insertion sites, DNA was sent to MiGS (Pittsburgh, PA) for Illumina sequencing following the standard workflow for library preparation and read trimming as described above. This workflow generated a total of 1,331,568 paired reads and 358,000,888bp (∼60× coverage) of sequence. Raw Illumina reads for the transposon libraries have been uploaded to the SRA at accession SRR25551084.

All genome sequences were assembled using unicycler v. as implemented in the Cyverse pipeline v0.4.8 (47, 48). For the *Ptt* genome, both short and long reads were included in the hybrid assembly while all other genome were assembled from short reads only.

### Bacteriocin induction and preparation

The method for bacteriocin induction and preparation has been previously described in detail(49). For the induction and isolation of bacteriocins, a single colony of *Ptt* picked, transferred to 2–3 mL of KB, and grown overnight on a shaking incubator at 220 rpm and 27 °C. The next day, the culture of the strain of interest was back-diluted 1:100 in 3 mL of KB and placed back on the shaking incubator for 3–4 h. After this incubation period, mitomycin C (MMC) was added to the 3 mL culture to a concentration of 0.5 µg mL^−1^ and incubated on the shaker overnight to allow for induction of bacteriocin production. The next day, 1–2 mL of the MMC induced cultures was centrifuged at 20,000 *g* for 5 min to form a pellet. Unless undergoing PEG precipitation (see below), the supernatant was then carefully transferred to a sterile microcentrifuge tube by filter sterilization using a 0.22 µm pore size filter. Supernatants were stored at 4 °C. For non-MMC-induced samples, overnight cultures were not back-diluted and were immediately filter-sterilized after the overnight incubation period.

### PEG precipitation

The method for PEG precipitation has been previously described in detail (49). Polyethylene glycol (PEG) precipitation is used for the separation and concentration of high molecular weight compounds. NaCl and PEG were added to the sterile supernatant at final concentrations of 1M and 10 %, respectively, and tubes were inverted repeatedly until dissolved. At this point tubers were incubated for either 1 h on ice or overnight at 4 °C. The samples are then centrifuged at 16,000 *g* for 30 min at 4 °C. After decanting the supernatant, the pellet was resuspended in a volume of 1/10 of the original supernatant volume of buffer (10 mM Tris, 10 mM MgSO_4_, pH 7.0). To remove any residual PEG and to kill any remaining bacteria, an equal volume of chloroform was added to the resuspended pellet and the tube was vortexed for 10–15 s, centrifuged for 20,000 *g* for 5 min, and then the upper aqueous layer was transferred to a clean microcentrifuge tube. This process was repeated for a total of two extractions until no white interface was seen between the aqueous and organic phase layers. Any residual chloroform was allowed to evaporate in a fume hood by leaving the tube uncapped for several (3–4) hours.

### Soft agar overlay

The method for the soft agar overlay assay used here has been previously described in detail(49). A single colony of the target strain was picked, transferred to 3 ml of KB liquid, and grown overnight on a shaking incubator at 220 rpm and 27 °C. The next day, the target strain culture was back-diluted 1:100 in 3 mL of KB and incubated for 3–4 h. Sterile 0.4 % water agar was melted and 3 mL was transferred to a clean culture tube and allowed to cool until warm to touch. An aliquot (100 µL) of the target strain culture was transferred to the melted water agar and vortexed to thoroughly mix. The water agar mixture was then poured onto a solid KB agar plate, swirled to cover the entire plate, and then allowed to cool for 15–20 min. Once solidified, 5 µL of the PEG-prepared supernatant was spotted onto the dish and allowed to dry. The plate was allowed to grow for 1–2 days and then the presence or absence of killing activity was recorded. All pictured soft overlay assays were repeated three times using independent bacteriocin inductions and overlays and results reported are representative of all replicates.

Two independent replicate experiments were carried out to measure the sizes of killing zones in soft overlay experiments. For each experiment, supernatant and PEG precipitated supernatant treatments were prepared as above from each strain with 5 µL of each spotted onto either *P. syringae* pv. syringae B728a (DAB2) or UB303R (DBL1887). For experiments using DAB2 as a target strain, there were 5 measurements total across the two independent replicate experiments. For experiments using UB303R as a target strain, there were 6 measurements total across the two independent replicate experiments. Overlay plates were scanned and area of the killing zones were measured using imageJ (50) (standardizing for 100mm diameter of petri dishes). Scans underlying raw data can be found at https://doi.org/10.6084/m9.figshare.24208449, while raw data and code to reconstruct Figure 1B can be found at https://doi.org/10.5281/zenodo.8384752.

### Transposon Mutant Isolation and Screening

For construction of transposon libraries, a single *E. coli* colony housing a single plasmid containing a modified Tn5 transposon was isolated from a previously constructed BarSeq library. This plasmid was conjugated with *Ptt* through standard methods, briefly, both strains were grown overnight in Lysogeny Broth (LB) media containing either rifampicin 50 ng/uL (*Ptt*) or kanamycin 30 ng/uL and diaminopimelic acid 200ng/uL (*E. coli*). Cells were pelleted and washed 2x in 10Mm MgCl2 with supernatant discarded after each wash, after which point the cells were mixed and washed in 1mL 10Mm MgCl2. The mixture was pelleted, with supernatant discarded, and the cells were resuspended in the remaining liquid. This mixture was plated on LB agar plates supplemented with 200 ng/uL DAP and incubated at 28oC. After 7 days, the conjugation mixture was resuspended and dilutions were plated on KB agar supplemented with 25 ng/uL kanamycin and 50ng/uL rifampicin.

Kanamycin resistant colonies arising from the conjugation were patched to KB agar containing 25ng/uL kanamycin, and after growth, picked to 200uL KB media in a 96 well plate and incubated at 28oC with shaking overnight. The next day, a drop of culture from each of the 96 wells was transferred to 100uL of KB media in a new 96 well plate and incubated for 4 hours at 28oC with shaking, after which point an additional 100uL of KB media containing 1 µg mL^−1^ MMC was added to the plate. The plate was then incubated overnight with shaking at 28oC. The next day, 3uL samples from each well of the plate was spotted onto a soft agar overlay plate prepared as described above except that for the overlay, 300uL of strain *Psy*B7828 was added to 5mL of 0.4% agar of the overlay strain and plated onto a 135mm petri dish containing KB media with streptomycin added to the top agar at a final concentration of 200ng/uL. Overlay plates were screened for lack of clearing zones, as indicative of the killing activity from *Ptt* against *Psy*B728a being eliminated.

### Creation of *Ptt* Strains Containing Directed Deletions

The creation of DBL1944 (a Rbp/ch deletion mutant of *Ptt*, referred to as *PttΔrbp/ch* throughout the text) was carried out using the Gateway cloning system with pMTN1907 as the destination vector. To create this mutant, a Gblock was ordered from IDT and BP clonase was used to recombine this dsDNA into plasmid pDONR207 to generate pDBL112. LR recombinase was then used to recombine a dsDNA segment into pMTN1907 to generate pDBL113and transformed into *E. coli* S17 lambda *pir*. As per previously described methods (CITATION), the *E. coli* strain containing pDBL113 was mixed with *Ptt* and incubated on Lysogeny broth plates (without antibiotic amendment) for 2 days. At this point, the conjugation mixture was resuspended and plated onto KB agar plates containing rifampicin 50 ng/uL, tetracycline 10 ng/uL, and nitrofurantoin 40 ng/uL. Kanamycin resistant *Ptt* isolates were streaked to pure single colonies on KB containing rifampicin and kanamycin at which point 8 independent colonies were picked to a 2mL KB culture, grown overnight, and then plated onto KB media containing 10% sucrose. Sucrose resistant colonies arising from this selection were patched to KB plates containing 10% sucrose plates and further patched to KB plates containing tetracycline. Sucrose resistant, tetracycline sensitive colonies were then screened for presence of the deletion by PCR. This Rbp/ch deletion was verified by whole genome sequencing, with raw sequencing reads available at accession TBD. DBL1945 (the aptatacin L1 deletion mutant of *Ptt*, referred to as *PttΔapt* throughout the text) was created by ordering a deletion vector from Twist Bioscience (San Francisco, CA USA) using pKM-18s (51) as a backbone vector to create pDBL114. This vector was conjugated into *Ptt* to create the deletion mutant using standard methods for pMTN1907 growth and selection (as described above, except that selection for integration fop pDBL114 into *Ptt* was performed and transconjugants were screened on KB agar plates containing rifampicin, nitrofurantoin, and kanamycin 25 ng/uL. For mutants arising from these procedures, a single colony was grown overnight at 27 °C with shaking in KB medium and frozen in 40% glycerol at −80 °C to. For all other *P. syringae* isolates, a single colony was grown overnight at 27 °C with shaking in KB medium supplemented with rifampicin 50 ng/uL.

### Expression of aptatacin in *E. coli*

A pET-28a(+) expression vector with the addition of the *apt* gene was obtained through Twist Bioscience and is referred to as pDBL115. The plasmid was transformed into *E. coli* strain BL21 (DE3) competent cells. Briefly, 1 µL of plasmid (50 ng) was added to 50 µL of competent cells and incubated on ice for 30 minutes. The cells were then heat shocked for 15 seconds at 42°C and immediately added to ice for 2 minutes. 500 µL of LB was added to the tube and incubated at 37°C for 1 hour. The transformation was plated on LB + kanamycin 25 µg/mL and incubated at 37°C. A single colony from the selection plate was grown in LB liquid + kanamycin 25 µg/mL overnight at 37°C. A midlog phase culture (0.3-0.6 OD_600_) was induced with IPTG for a final concentration of 0.33 mM IPTG and incubated overnight at 22°C with shaking. Cells were spun down at 8,000 x g for 2 minutes and supernatant decanted. Pellet was resuspended in 500 µL of lysis buffer (20 mM Tris-HCl, 500 mM NaCl) and lysed using sonication. The lysed solution was spun down and supernatant collected for overlay spotting.

## ACKNOWLEDGEMENTS

This work was supported by a grant from the National Science Foundation (NSF) (IOS 1856556) to DAB. We thank Meara Clark and Kevin Hockett for extensive help with overlay experiments.

